# Spatial, Quantitative and Functional Deconstruction of Virus and Host Protein Interactions Inside Intact Cytomegalovirus Particles

**DOI:** 10.1101/2022.05.02.490278

**Authors:** Boris Bogdanow, Iris Gruska, Lars Mühlberg, Jonas Protze, Svea Hohensee, Barbara Vetter, Martin Lehmann, Lüder Wiebusch, Fan Liu

**Author notes:** correspondence; fliu@fmp-berlin. co-senior authors (Lüder Wiebusch, Fan Liu).

## Abstract

Herpesviruses assemble large enveloped particles that are difficult to characterize structurally due to their size, fragility and complex proteome with partially amorphous nature. Here we use cross-linking mass spectrometry and quantitative proteomics to derive a spatially resolved interactome map of intact human cytomegalovirus virions. This enabled the *de novo* allocation of 32 viral proteins into four spatially resolved virion layers, each organized by a dominant viral scaffold protein. The viral protein UL32 engages with all layers in an N-to-C-terminal radial orientation bridging nucleocapsid to viral membrane. In addition, we observed the layer-specific recruitment of 82 host proteins, a subset of which are constitutively and selectively incorporated via specific host-virus interactions. We uncover how the recruitment of PP1 phosphatase and 14-3-3 proteins by UL32 affects early and late steps during viral biogenesis. Collectively, this study provides global structural insights into the native configuration of virus and host protein interactions inside herpesvirus particles.

## MAIN

The structured assembly of infectious particles, called virions, is fundamental for virus transmission among cells and organisms. Virions contain the viral nucleic acid genome enclosed in a capsid protein shell, and a number of co-packaged proteins, which facilitate the infection process and the onset of viral gene expression. Herpesviruses, a family of double-stranded DNA viruses, assemble particularly large and complex particles, accommodating many different proteins that are delivered to the host cell upon infection.

The capacity of herpesvirions to incorporate a large set of proteins is enabled by their typical multilayered architecture ^1^. The outer lipid envelope harbors various viral glycoproteins required for host cell receptor binding and membrane fusion ^2^. The space between the envelope and the central, icosahedral nucleocapsid is filled with a proteinaceous matrix, the tegument. While individual tegument proteins have been allocated to distinct inner- and outer sub-layers based on biochemical ^3^ and microscopic data ^4, 5^, the details of tegument protein organization are not understood Herpesvirus particles also incorporate numerous host proteins, but very few of these events have been functionally or mechanistically characterized ^6, 7^.

Recent cryoEM studies of herpesvirions have revealed substructures of the nucleocapsid ^8, 9^, the portal ^10, 11^ and several glycoprotein complexes ^2, 12, 13^. In addition, previous studies only provided insight into the overall protein composition of herpesvirions ^14^, but a systematic characterization of the spatial coordination and interactions of virus and host proteins within virions is lacking.

Here, we use cross-linking mass spectrometry (XL-MS) to build a virion-wide proximity map of 32 viral and 82 host proteins in intact extracellular virions of human cytomegalovirus (HCMV), the largest human herpesvirus. The data enable a *de novo* allocation of host and virus proteins and their protein-protein interactions (PPIs) to virion layers, providing insights into the organization of the tegument sublayers. We find that the viral protein UL32 (also known as pp150) acts as a dominant scaffold, engages in PPIs across the particle and mediates the recruitment of host proteins, such as 14-3-3 proteins and protein phosphatase 1 (PP1). PP1 antagonizes 14-3-3 binding to UL32 and is required for efficient start of viral gene expression and production of viral progeny. Thus, by charting the proteome organization within native herpesvirions, we provide a basis for the structural and functional understanding of crucial PPIs.

## RESULTS

### A protein-protein interaction network of the intact HCMV particle

XL-MS allows capturing protein contacts from native environments ^15^ such as organelles ^16–20^. We reasoned that XL-MS is also suitable for gaining global insights into PPI networks of large and structurally heterogeneous herpesviral particles.

Therefore, we isolated extracellular particles from infectious cell culture supernatant and cross-linked them with disuccinimidyl sulfoxide (DSSO), which connects lysines from proteins in close spatial proximity (Fig. 1a). After purification of the cross-linked virions using tartrate-glycerol density gradient centrifugation, protein extraction and tryptic digestion, the cross-linked peptides were identified by liquid chromatography mass spectrometry (LC-MS).

**Fig. 1.**
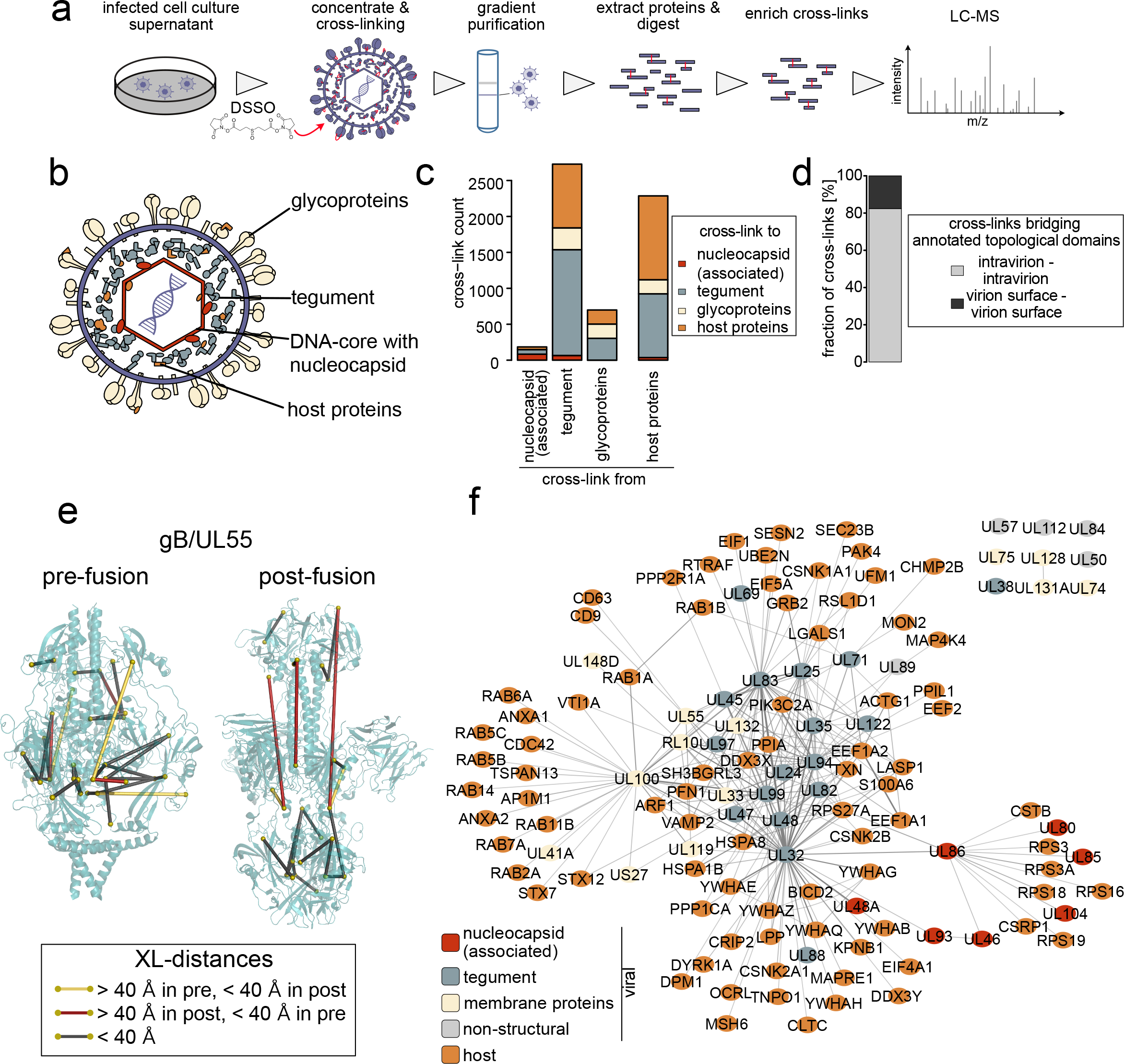
The spatial proteome of intact cytomegalovirus virions. **(a)** Workflow for the XL-MS analysis of intact HCMV particles. The experiment was performed in n=2 biological replicates. **(b)** Schematic depiction of the HCMV virion layers. **(c)** Number of cross-links among viral proteins with known virion layer localization and between viral and host proteins. **(d)** Proportion of cross-links within extra-virion and intra-virion domains of viral glycoproteins or intra-virion resident proteins. No cross-links have been observed linking extra-virion domains of viral glycoproteins to their intra-virion domains or to intra-virion resident proteins. **(e)** Cross-link mapping onto structural models of UL55 in pre- and post-fusion conformations. See also Extended Data Figs. 1a-c. **(f)** Network of PPIs inside HCMV particles. The line width (edges) scales with the number of identified cross-links between interaction partners.

We identified 9,643 unique lysine-lysine connections at a 1% false discovery rate (FDR) (Supplementary Table 1). First, we asked whether these data correctly captured the overall spatial organization of the virion (Fig. 1b) and counted the cross-links within and between proteins having previously reported virion layer localization (Fig. 1c). We found most cross-links within the tegument, followed by the envelope and the nucleocapsid. Importantly, we did not observe any cross-links connecting viral membrane proteins to viral nucleocapsid proteins, confirming that the cross-links reflect the known layered virion architecture. Furthermore, we did not identify cross-links connecting the interior of the virion to extra-virion domains of viral glycoproteins (Fig. 1d), providing confidence that the viral particles were not damaged prior to cross-linking. Host proteins contributed to cross-links in all virion layers, confirming their association with herpesvirus particles^21–24^.

Next, we investigated whether our XL-MS data are in agreement with published HCMV protein structures. Considering that two protein side chains can only be cross-linked if their mutual distance is small enough to be bridged by the cross-linker, the linear distance between the cross-linked lysines should be below 40 ngstrom (L). To analyze whether this constraint is met, we mapped the cross-links on structural models of the homotrimeric viral glycoprotein UL55 (also know as gB) in its pre- and post-fusion state^13^ (Fig. 1e, Extended Data Figs. 1a-b). We identified 16 cross-links that agree with both the pre- and the postfusion structures. In addition, 3 cross-links could only be explained by the pre-fusion and 2 cross-links could only be explained by the post-fusion structure (Extended Data Fig. 1c). These conformation-specific cross-links confirm the previous finding that UL55 exists in different functional states ^25^ and demonstrate that we capture biologically relevant structural states of viral proteins.

We then set out to build an XL-based map of intra-virion PPIs and applied further filtering criteria on our list of cross-links to increase confidence (Extended Data Fig. 1d). This resulted in a reduced list of 2,248 cross-links supporting 262 PPIs (143 virus-host, 20 host-host, 99 virus-virus PPIs) with 82% overlap between two biological replicates (Supplementary Table 2, Extended Data Fig. 1e, Fig. 1f). The highly connected center of the network contains tegument proteins that are frequently linked to each other and to neighboring virion layers, in line with the characterization of the tegument as a highly interconnected module bridging nucleocapsid and viral envelope ^26^. Viral membrane proteins as well as nucleocapsid proteins cluster independently and are in general much less connected.

### *De novo* allocation of viral proteins into distinct virion layers

While the localization and structure of the viral transmembrane and nucleocapsid proteins are well established, the spatial organization of the viral tegument is largely elusive. We hypothesized that the proximity-based nature of our data should enable the *de novo* allocation of viral tegument proteins to distinct layers within the particle.

To this end, we calculated ‘PPI specificity’ values for all viral PPIs as the ratio of cross-links between two interaction partners (“interactor 1” and “interactor 2”) to the total number of cross-links of the “interactor 1” proteins (Fig. 2a). We treated each viral protein as interactor 1 (columns in Fig. 2a) and as interactor 2 (rows in Fig. 2a) and performed hierarchical clustering. This yielded two main clusters that can both be separated into two smaller clusters. As expected, nucleocapsid and viral transmembrane proteins separated into distinct main clusters (clusters 1 and 2 in Fig. 2a) and further contained two subclusters, supporting the view that the four subclusters represent the four architectural virion layers.

**Fig. 2.**
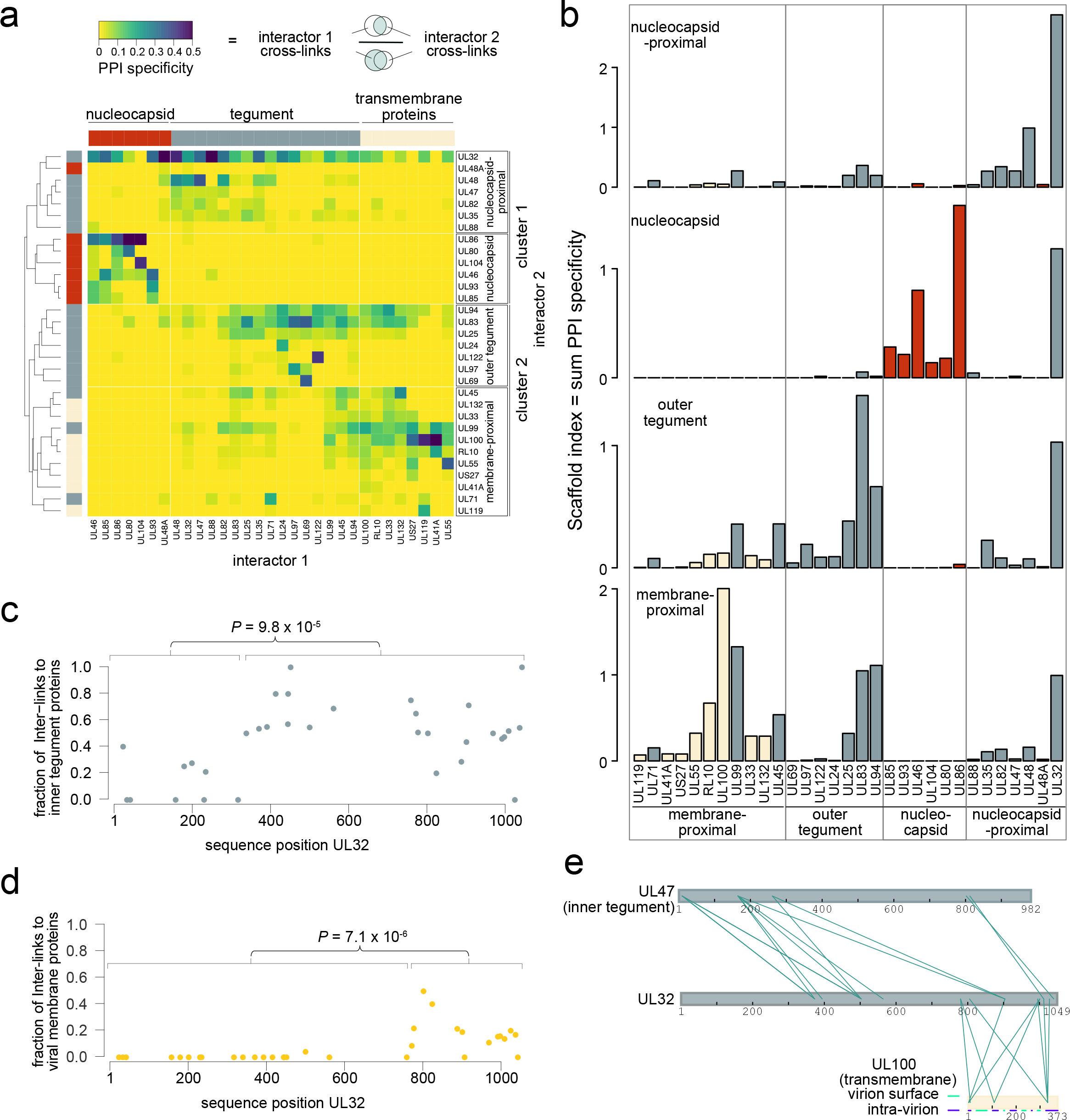
Spatial arrangement of viral proteins. **(a)** Heatmap of PPI specificity values of all viral proteins with at least 9 identified crosslinks. PPI specificity values were calculated for each PPI by dividing the number of cross-links supporting a PPI by the total number of crosslinks of interaction partner (‘interactor 1’). Interactor 2 proteins (rows) were clustered using 1-Pearson’s correlation distance and complete linkage. Interactor 1 proteins (columns) were manually annotated based on prior knowledge. **(b)** By summing up the PPI specificity values of interactor 2 proteins across all PPIs of the respective subcluster, a scaffold index was calculated and plotted using the same order of proteins as in the columns of panel A. Color schemes indicate the membership of proteins to nucleocapsid, tegument or viral envelope (transmembrane proteins). **(c-d)** For each individual lysine in UL32, the fraction of cross-links in UL32 linking to other viral inner tegument proteins **(c)** or viral membrane proteins **(d)** is plotted. *P*-values are based on two-sided wilcoxon rank sum tests comparing the indicated lysines within the UL32 primary sequence. **(e)** Cross-link network between UL32 and selected proteins of the inner tegument (UL47) and viral envelope (UL100). The membrane topology of UL100 is indicated by blue and green lines.

Focusing on the main cluster 1, we observed that nucleocapsid and tegument proteins form distinct subclusters. Only the smallest capsid protein UL48A, for which we detected relatively few cross-links, co-clusters with the tegument group of cluster 1. Besides UL48A, this subcluster contained typical inner tegument proteins like UL32 and UL48, that are characterized by their tight association to the nucleocapsid ^3^ as well as the UL48-associated protein UL47 ^27^,UL82 (also known as pp71), the UL82 interactors UL35 ^28^ and UL88, which are important for recruiting UL47, UL48 and UL82 into virions ^29^. As UL82 and UL35 proteins traffic independently from the nucleocapsid upon infection ^30, 31^, their allocation to the nucleocapsid-proximal tegument layer indicates that proximity to the nucleocapsid does not necessarily reflect tight attachment upon cell entry.

The second main cluster (cluster 2) contained two subclusters, one with viral transmembrane proteins and tegument proteins and the other with tegument proteins only. The first subcluster contained tegument proteins UL71, UL99 (also known as pp28) and UL45, indicating their membrane association, consistent with previous reports ^32, 33^. The second distinct subcluster represents the outer, non-membrane bound tegument. Its composition fits previous observations showing that the UL83 (also known as pp65) protein is important for incorporating UL69, UL97 and UL25 into mature particles ^22, 34^. Thus, based on our cluster analysis, we define at protein-level resolution a distinct nucleocapsid layer and three tegument sublayers, including a nucleocapsid-proximal inner tegument, an outer tegument and a membrane-associated tegument. The identification of three separately organized tegument sub-structures points to a potential hierarchy in the herpesvirus tegumentation process.

After establishing the layer-specific organization of viral proteins, we asked which of these proteins are most important for the overall organization of PPIs. We reasoned that such proteins would act as scaffolds that specifically recruit other proteins. To find these, we summed up the ‘PPI specificity’ values of each viral protein in the respective subcluster (Fig. 2b), which revealed several main scaffolds within the individual layers: UL86 (also known as major capsid protein MCP) for the nucleocapsid, UL32 for the nucleocapsid-proximal tegument, UL83 for the outer tegument and UL100 (also known as gM) for the membrane-proximal proteins. Importantly, UL32 contributed to scaffolding in all layers and had the overall highest scaffold index, indicating that it serves as an organizational hub.

Despite its allocation to the nucleocapsid-proximal inner tegument, UL32 cross-linked with proteins from all layers of the virion (Fig. 2a, top row): Nucleocapsid (e.g. UL86), inner tegument (e.g. UL48, UL47), outer tegument (e.g. UL83) as well as viral glycoproteins (e.g. UL55, UL100). Considering that UL32 is anchored with its N-terminal domain at the nucleocapsid ^35^, we reasoned that the disordered C-terminal domain (AA 303-1049) may engage in interactions with the other layers. Analyzing the cross-linking partners of each UL32 lysine residue shows that residues 340-1049 engage more frequently with other inner tegument proteins (Fig. 2c). Likewise, a region in UL32 (residue 775-end) associates with viral membrane proteins (Fig. 2d). This indicates that a central region within UL32 locates to the tegument, whereas the more C-terminal distal region associates with both the tegument and the viral membrane. This domain-specific interaction pattern is further exemplified by its domain selective cross-linking to an inner tegument protein UL47 and a transmembrane protein UL100 (Fig. 2e). Thus, UL32 engages with all layers of the particle in an N-to-C terminal radial orientation bridging nucleocapsid to viral membrane.

### Layer-specific organization of the host proteins inside the virion

After establishing the organization of viral proteins within the particle, we turned to the 82 incorporated host proteins. 79 of these host proteins have been observed in previous proteomic studies as associated with purified virions using standard proteomics ^21–24^ (Extended Data Figs. 2a-b). First, we determined their location within the layered herpesvirion by analyzing cross-links between individual host proteins and viral proteins of the specific virion layers, resolving host protein localization to these layers (Fig. 3a). For example, ribosomal 40S proteins (RPS3A, RPS18, RPS16, RPS3, RPS19) specifically cross-linked with the viral nucleocapsid at four distinct lysine residues on the UL86 protein (Extended Data Fig. 2c). Mapping these cross-links on a hexon structure revealed that ribosomal proteins associate with the nucleocapsid interior (Extended Data Fig. 2d), which is predominantly positively charged as it has to accommodate viral DNA. We reason that electrostatic interactions between negatively charged ribosomes and the positively charged nucleocapsid interior likely explain this association.

**Fig. 3.**
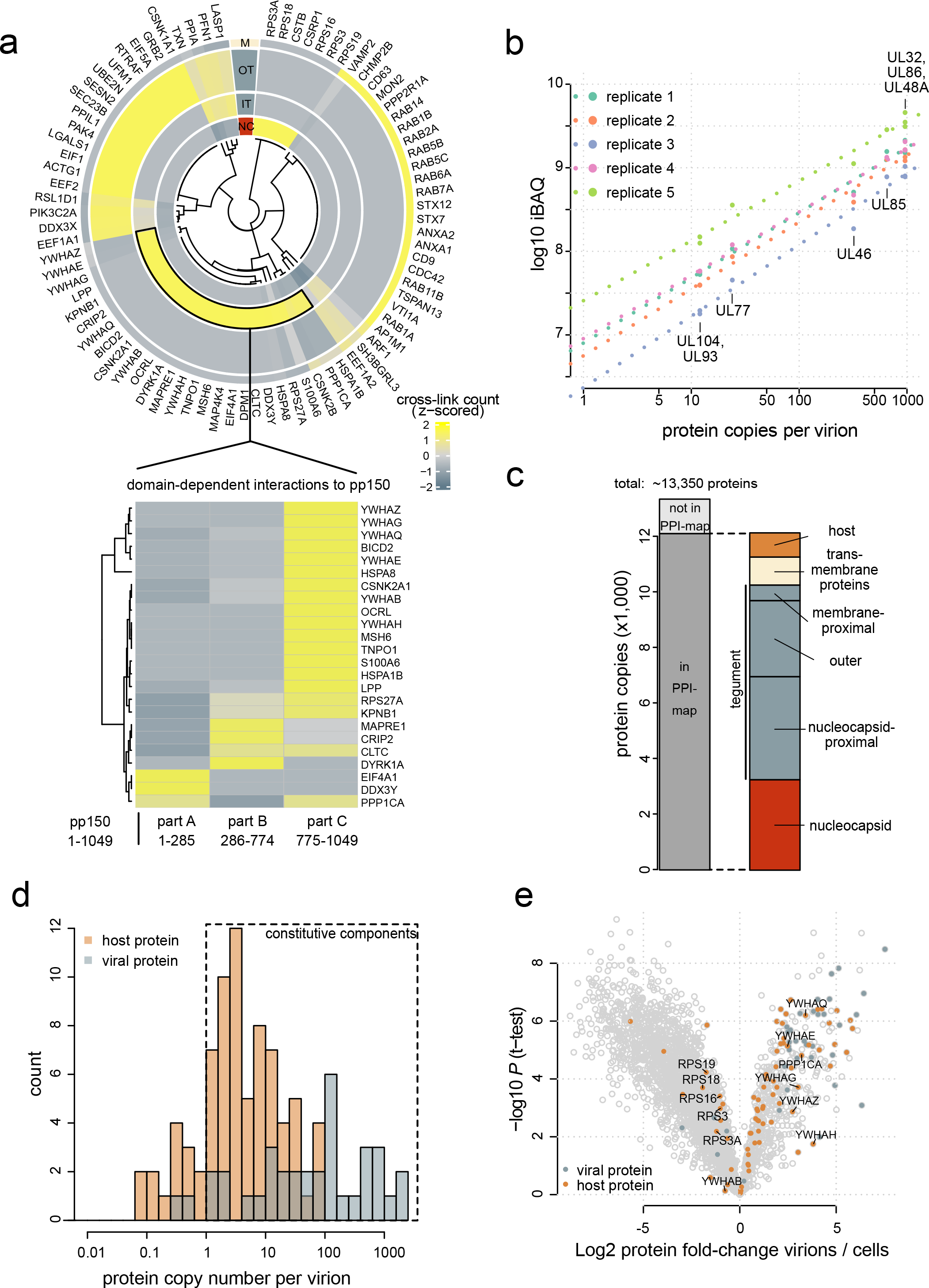
Abundant and layer-specific incorporation of host proteins. **(a, top)** Heatmap depiction of z-scored cross-link counts across virion layers. Viral proteins with established localization were used as markers for sub-virion localization. NC: nucleocapsid, IT: inner tegument (nucleocapsid-proximal tegument), OT: outer tegument, M: membrane, including viral transmembrane proteins and membrane-proximal tegument. **(a, bottom)** Z-scored cross-link counts of inner tegument localized host proteins to different parts of UL32. **(b)** Intensity-based absolute quantification of protein copy numbers using the copy numbers of nucleocapsid associated proteins as standard (n=2 biological replicates, with technical duplicates or triplicates). The offset between the replicates is caused by differences in the amount of input material but the similar slopes demonstrate reproducibility of the quantitative data. **(c)** Protein copies represented in the PPI map and their sub-virion localization. **(d)** Histogram of copy numbers from host and viral proteins. Proteins with more than one copy number on average are classified as likely constitutive. **(e)** Relative quantification of proteins comparing virion lysates to cell lysates. Proteins included in the PPI map are highlighted. See Extended Data Fig. 4a for experimental design. Means of fold-change differences and t-test P values are based on n=4 biological replicates.

The inner tegument incorporated a Lim-domain containing protein (LPP), heat-shock proteins (HSPA8, HSPA1), a cargo-adaptor (BICD2) and clathrin (CLTC). Additionally, we found various phospho-regulatory proteins in the inner tegument, such as 14-3-3 proteins (YWHAx, with x being either B, E, G, H, Q or Z), kinases (DYRK1A, casein kinases CSNK2B, CSNK2A1) and PP1 (PPP1CA). Most of the inner tegument-resident host proteins were cross-linked to UL32 predominantly to the region close to its C-terminus (Fig. 3a, bottom).

The outer tegument contained proteins involved in innate immunity (DDX3X), translation factors (EEF2, EEF1A1, EIF5A, EIF1), chaperones (PPIA, PPIL1) and cytoskeletal proteins (ACTG1, PFN1, LASP1). At the viral envelope we observed host proteins related to exo- and endocytosis and membrane trafficking, such as RAB-proteins (RAB1A/B, RAB2A, RAB5B/C, RAB6A, RAB11B, RAB14), adaptor proteins (AP1M1), snare proteins (VAMP2, VTI1A), annexins (ANXA1, ANXA2), syntaxins (STX7, STX12) and the tetraspanins TSPAN13, CD63, CD9. Interestingly, the latter two tetraspanins were found to be cross-linked at the outside of the particle, associating with the UL55 virion surface domain (Extended Data Fig. 3e).

Collectively, this spatially resolved map of host proteins within virions indicates that host proteins are localized to different layers, reflecting the physical encounters of host proteins during different stages of virion assembly. Furthermore, UL32 and in particular its C-terminal region engages in many interactions with host proteins, consistent with its scaffolding role for viral proteins.

### Quantitative assessment of host protein recruitment

After analyzing the sub-virion localizations of viral and host proteins, we aimed to characterize the quantitative dimension of their incorporation into the viral particle. To acquire an accurate protein inventory of the particle, we calculated copy numbers of host and viral proteins (Fig. 3b, Supplementary Table 3) and found that HCMV virions contain on average ∼13,350 protein copies (Fig. 3c). Considering the spherical diameter of 200 nm, this results in a virion protein density of about 3.2*10^6^ proteins/ cm^3^, similar to density estimates for mammalian and prokaryotic cells ^36^. 91% of the copies belong to proteins that are contained in our virion network, indicating that we obtained a comprehensive spatial characterization of an average particle. Importantly, the vast majority of these proteins are incorporated with more than 1 copy per virion, indicating that these proteins are, on average, constitutive components (Fig. 3d).

We next asked whether host proteins are specifically targeted to virions, which may indicate functional relevance. We hypothesized that host proteins are incorporated either non-selectively, especially if they are highly abundant in the host cell, or based on specific interactions with viral proteins promoting their recruitment. These scenarios can be distinguished by comparing the relative levels of host proteins in the virion and in cells, where increased protein levels in the virion relative to the cell are indicative of active recruitment. To address these options, we harvested infectious cell culture supernatant for virion purification and the infected cells in parallel, and analyzed both samples by quantitative proteomics (Extended Data Fig. 3a, Supplementary Table 4) with overall good reproducibility between replicates (Extended Data Fig. 3b). While most host and viral proteins contained in our spatial virion map were significantly enriched in virion lysates, several proteins were not enriched or even modestly depleted in virion preparations (Fig. 3e). Comparing enrichment levels to copy numbers (Extended Data Fig. 3c), we found that some proteins, such as heat-shock proteins, ribosomal proteins and translation factors were constitutively incorporated but only to levels that reflect their cellular abundances indicating that they are non-specifically packaged into virions.

In contrast, several low-abundant cellular proteins such as the cargo transporter adaptor BICD2, the tetraspanin TSPAN13 and the phosphodiesterase OCRL were strongly enriched in virions. While these proteins are not among the most abundant components of the virion, they exclusively cross-linked to one viral protein (Extended Data Figs. 3d-e), supporting the idea that they are actively recruited to the virion through specific PPIs.

### UL32 C-terminal domain recruits 14-3-3 and PP1 via specific binding motifs

We then selected PP1 and 14-3-3 proteins for further investigation because they are among the most abundant hot proteins that are selectively recruited into virions according to our quantitative data (Extended Data Fig. 3c). They cross-linked to the membrane-proximal tegument protein UL99 and, more frequently, to UL32 (Fig. 4a). We validated the incorporation of 14-3-3 and PP1 into the tegument and their association with UL32 by double-immunogold labeling and electron microscopy analysis of intracellular HCMV particles (Fig. 4b, Extended Data Fig. 4a-b).

**Fig. 4.**
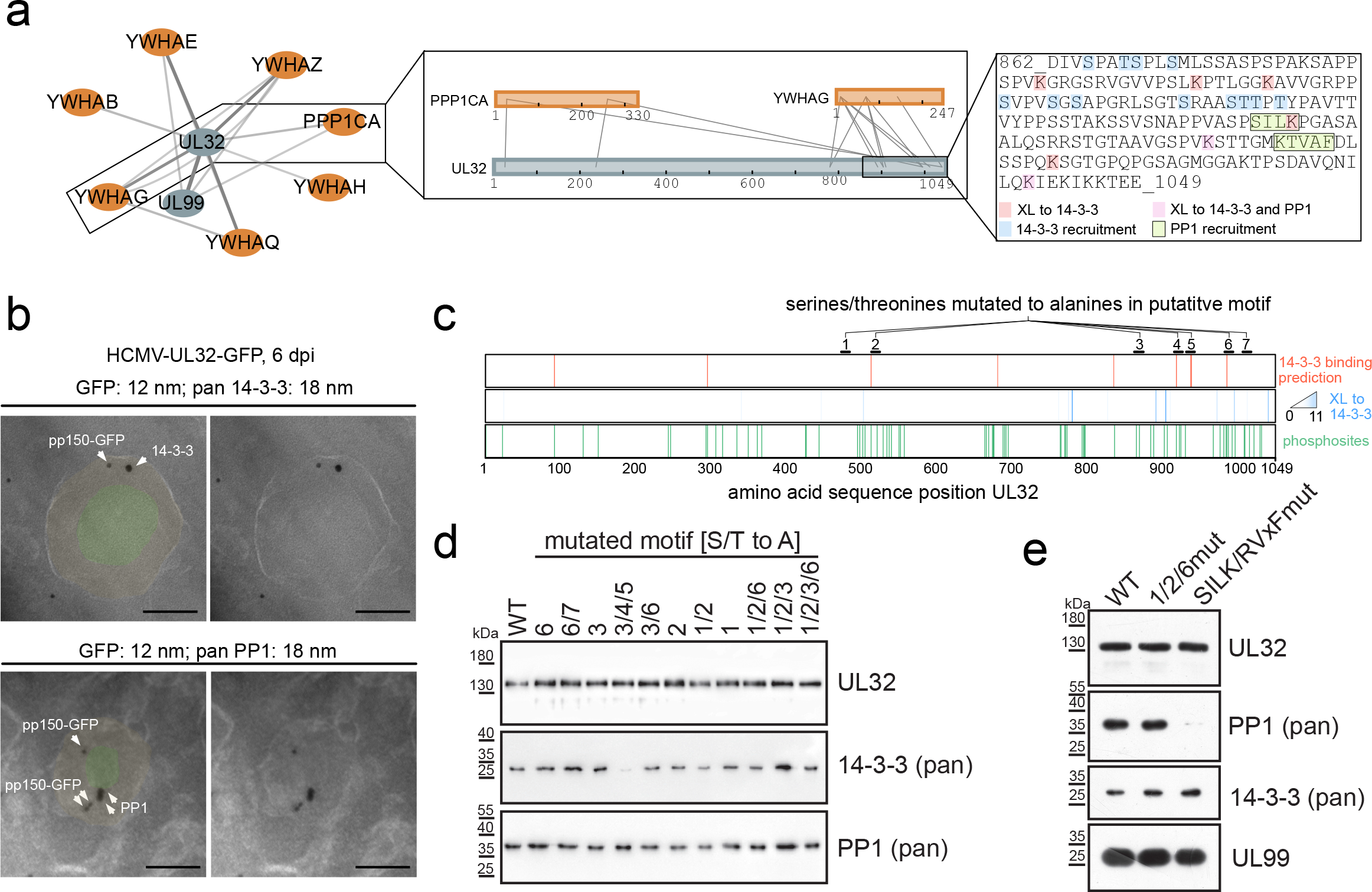
Nearby short linear motifs in UL32 recruit 14-3-3 and PP1 proteins into virus particles. **(a)** PPI network including interactors of PP1 (PPP1CA) and 14-3-3 (YWHA*) proteins with insets showing cross-links involving UL32, 14-3-3 protein gamma (YWHAG), PP1 and the UL32 sequence containing 14-3-3 and PP1 recruitment sites. The line width scales with the number of identified cross-links. Host and viral proteins are highlighted orange and grey, respectively. **(b)** EM images of intracellular virions from HCMV-UL32-GFP infected fibroblasts (MOI=5). Ultrathin sections were stained with immunogold against GFP (12 nm gold) and either pan 14-3-3 or PP1 (18 nm gold), as indicated. Approximate areas of tegument and nucleocapsid layers were manually highlighted in light orange and light green. Scale bars 100 nm. **(c)** Mutational approach for identifying recruitment sites for 14-3-3 on UL32. Motif predictions above score 0.6 (top bar), cross-link positions to 14-3-3 (middle bar), identified phosphorylation sites (localization probability > 0.75 in n=2 biological replicates) in virions (bottom bar). **(d,e)** Purified virions from recombinant mutant viruses harboring alanine substitutions in the designated motifs were assessed for protein levels by immunoblotting. Representative experiments of n=2 biological replicates are shown.

We then hypothesized that PP1 and 14-3-3 proteins are recruited via interaction motifs in UL32. Dimeric 14-3-3 proteins typically bind to phosphorylated serines or threonines within motifs that can be predicted by machine learning based on the local sequence context ^37^. Combining these predictions with information from our cross-link data, measured phosphorylation sites of UL32 in virions (Supplementary Table 5, Fig. 4c), we hypothesized 7 potential motifs as 14-3-3 binding sites. We then mutated serines and threonines within these 7 potential motifs to alanines in different combinations in the viral genome. Combined mutation of three motifs abolished 14-3-3 incorporation into the particle (Fig. 4d). Comparison of the virion proteome of this mutant to wild-type (WT) viruses using quantitative proteomics based on Stable Isotope Labeling by Amino Acids in Cell Culture (SILAC) further indicated that 14-3-3 proteins were selectively lost from mutant virions without strongly affecting other host or viral proteins (Extended Data Figs. 4c-e).

Next, we investigated the recruitment of PP1 into HCMV particles, a characteristic of HCMV ^38^ with unknown mechanistic basis and biological significance. PP1 has several surface grooves that bind to interactors containing for instance RVxF and SILK motifs ^39^. In addition, UL32 harbors these motifs in its C-terminal 100 amino acids (Fig. 4a) and cross-links in close spatial proximity to the RVxF binding groove (Extended Data Fig. 5a). Again, we mutated critical amino acid residues to alanine (SILK/RVxFmut) within these motifs, purified the virions and assessed the levels of PP1. Incorporation of PP1 was substantially reduced with the mutant virus (Fig. 4e), which was confirmed by SILAC-based analysis of the whole proteome of mutant and WT virions (Extended Data Fig. 5b). The abundances of most other proteins were not altered in the mutant virions.

However, PP1 depletion correlated with depletion of DYRK1A, another specific cross-linking partner of UL32, (Extended Data Fig. 5c) and DYRK1A-interactors, such as the APC/C complex subunits CDC23 and CDC16 ^40^, and DCAF7 ^41^ . A phosphopeptide analysis of these SILAC-labeled mutant and WT virions (Extended Data Fig. 5d) showed that PP1 depletion correlated with significantly higher abundances of phosphopeptides derived from UL32 compared to other viral or host proteins, indicating that PP1 preferentially dephosphorylates UL32. Thus, PP1 is recruited to the particle via SILK/RVXF motifs that are in proximity to 14-3-3 recruitment motifs, confirming that host protein incorporation into viral particles is mediated by specific host-virus interactions.

### PP1-recruitment regulates early and late events during HCMV biogenesis

To test the functional relevance of the UL32-controlled recruitment of PP1 to viral particles, we first investigated the ability of the UL32-PP1-binding deficient mutant virus (SILK/RVXFmut) to replicate in cell culture. Compared to the parental WT virus, the mutant virus produced up to 20-fold fewer viral progeny (Fig. 5a, Extended Data Fig. 6a), supporting that recruitment of PP1 to UL32 is functionally important for efficient production of viral progeny.

**Fig. 5.**
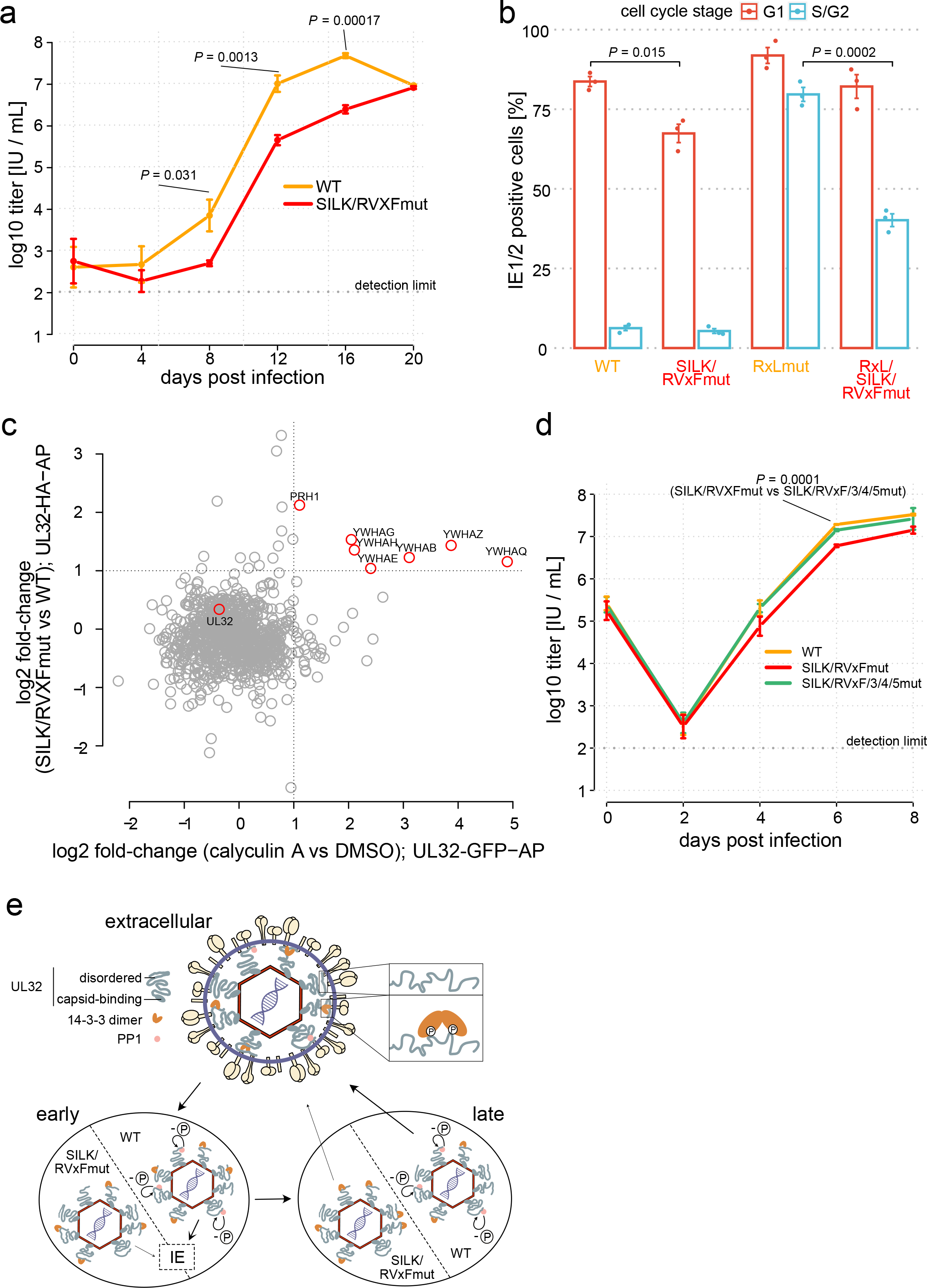
PP1 recruitment controls early and late events during the HCMV biogenesis. **(a,d)** Growth curve of mutant and wildtype virus on primary lung fibroblasts (PLFs) (panel a: MOI=0.05, panel e: MOI=5). Means (center of the error bars) and standard deviations of the mean of *n* = L 3 biological replicates are depicted. Unpaired two-sided *t*-tests without multiple hypothesis correction were performed at the indicated time points. **(b)** Flow cytometry analysis of Immediate Early (IE) protein levels as a function of the cell cycle stage. PLFs were infected with the indicated recombinant viruses (MOI = 5, 6 h post infection). The percentage of IE1/2-positive cells in G1 or S/G2 compartments is given with mean (center of the error bars) and standard deviations (error bars) of n=3 biological replicates. Unpaired two-sided t-tests were performed comparing the fraction IE1/2 positive cells between the indicated viruses and cell cycle compartments. Representative dot-plots of one replicate are shown in Extended Data Fig. 6b. **(c)** Comparison of differential interaction partners to UL32 upon phosphatase inhibition (x-axis) or genetic disruption of the SILK/RVxF motifs (y-axis). See Extended Data Fig.7 for volcano plots and experimental design. **(e)** Location of 14-3-3 and PP1 inside virions and their role during early and late stages of HCMV infection. WT viruses, able to recruit PP1, dephosphorylate (-P) UL32 efficiently (bold arrows), start viral gene expression (IE) and produce infectious progeny. PP1-binding deficient RVxF/SILKmut viruses are impaired (light arrows) in the start of IE-gene expression and production of novel progeny.

While virion-delivered PP1 and UL32 enter the cell as part of the tegument, new copies of UL32 are produced in the infected cell only during later infection cycles. PP1-UL32 binding might thus be important during (i) events directly after entry before viral gene expression has started, (ii) during late stages when new copies of UL32 are produced or (iii) both.

We first focused on early events and assessed the production of the very first viral antigens (IE1/2-proteins). As the onset of HCMV gene expression is blocked during S/G2 phases of the cell cycle ^42^, we analyzed IE1/2-levels throughout the cell cycle by FACS analysis (Fig. 5b, Extended Data Fig. 6b). During susceptible G1, the WT virus accumulated IE-proteins in 84% of cells, which dropped to 62% with PP1-binding deficient SILK/RVxFmut virus. To assess the contribution of PP1 during the non-susceptible stage S/G2, we analyzed the PP1-binding site mutations in the genetic background of a virus able to start IE1/2-gene expression in S/G2 ^43^. Integrating the PP1-binding site mutations into this backbone (RxLmut) led to a decrease in the fraction of cells expressing IE1/2-proteins from 76% to 36% during S/G2. Together, this implicates UL32-recruited PP1 phosphatase as crucial for the onset of viral gene expression.

We then explored the role of the PP1-UL32 association during later stages of the infection cycle and asked whether PP1 modulates the interactions of UL32 with other proteins. To this end, we performed two interaction proteomics experiments: First, comparing the binders of WT-UL32 to the SILK/RVxFmut-UL32 (Extended Data Fig. 7a-b) and second, comparing the binders of WT-UL32 under treatment with phosphatase inhibitor calyculin A to a control (Extended Data Fig. 7c-d). We observed that, both the genetic disruption of PP1-UL32 binding and chemical inhibition of phosphatase activity leads to increased presence of 14-3-3 on UL32 (Fig. 5c), indicating that the presence of active PP1 phosphatase limits the association of UL32 with 14-3-3 proteins. To assess the functional relevance of this, we created a virus in which both the PP1 and the 14-3-3 binding sites were mutated (SILK/RVxF/3/4/5mut). Comparing this mutant to the PP1-binding deficient but 14-3-3-binding-competent SILK/RVxFmut virus showed that abolishing 14-3-3 recruitment rescued viral multiplication to almost WT levels (Fig. 5d). Thus, the phosphatase protects HCMV from over-loading 14-3-3 proteins onto the inner tegument, implying a perturbing or antiviral role for 14-3-3 proteins antagonizing efficient production of viral progeny.

## DISCUSSION

We integrated XL-MS and quantitative proteomics with molecular virology to provide detailed insights into the stoichiometry, architecture and domain-level PPIs of an infectious HCMV particle. Our study addresses long-standing questions regarding the organization of herpesvirions, as it i) enabled *de novo* reconstruction of the layered virion architecture including the spatial organization of the structurally complex viral tegument, ii) allowed comprehensive mapping of host-virus PPIs and their interaction contacts in the native configuration of an intact virion and iii) demonstrated the biological significance of recruited host proteins.

Based on our unbiased clustering of the viral tegument, we cataloged tegument proteins into three sub-layers: a nucleocapsid-proximal inner, an outer and a membrane-proximal tegument. Remarkably, we observed that the different tegument sub-layers are bridged by UL32. While its N-terminal domain is tightly capsid-bound ^35^, its C-terminal 250-300 amino acids associated with viral envelope proteins. The capsid-to-membrane tethering architecture of UL32 may be critical for the structural integrity of the virion and may also explain why UL32 is important during cytosolic maturation events ^44, 45^. The UL32 C-terminal region was found cross-linked not only to envelope but also to tegument proteins, suggesting that XL-MS captured different co-existing functional UL32 states as also observed for UL55 ^25^. In agreement with its disordered state ^43^, the UL32 C-terminus is thus likely to adopt multiple structural arrangements, of which only a fraction (∼20%) makes membrane contact.

This structural flexibility of UL32, paired with its high abundance and considerable sequence length, allows it to act as the dominant scaffold in the virion, organizing many interactions with other viral and cellular proteins. Importantly, UL32 associated with a variety of phospho-regulatory proteins and recruited 14-3-3 and PP1 via nearby binding sites in its C-terminus. Recruitment of PP1 to UL32 modulates the phosphorylation status of UL32, is important for high-titer replication of HCMV and positively regulates the onset of viral gene expression. At the late stage of infection, PP1 functionally and biochemically antagonizes the binding of 14-3-3 to UL32, indicating that HCMV recruits PP1 to limit antiviral or perturbing effects of 14-3-3. It is conceivable that overloading of the rigid 14-3-3 proteins might structurally restrain the flexible UL32 C-terminus and impair efficient virion assembly. These findings illustrate how host-virus interactions found within viral particles are relevant for early and late events of the replicative cycle.

Besides 14-3-3 and PP1, we found many other host proteins associated with the virion, ranging from the nucleocapsid interior to the virion surface. Such host proteins are constitutively incorporated based on two main routes. First, specific host-virus interactions lead to the selective enrichment of host proteins irrespective of their cellular abundance. One example is the cargo-adapter protein BICD2, which is known to facilitate trafficking and nuclear import of HIV-1 genomes ^46^. We found it specifically associated with UL32, suggesting that it may allow movement of the incoming HCMV-nucleocapsid along microtubuli, in analogy to cellular kinesin in HSV-1 virions ^6^. Second, non- or low-specific host-virus interactions lead to incorporation of highly abundant cellular proteins that are not or only modestly enriched. Among those are ribosomal proteins that we found associated with the DNA-accommodating side of the capsid. This association is sub-stoichiometric and consistently, ribosomal proteins are not overrepresented in virions compared to cells.

The understanding of the HCMV virion architecture has substantially improved through recently reported atomic structures of the nucleocapsid ^8^ and viral glycoproteins ^13, 47, 48^. This study further expands the understanding of herpesvirions by providing global structural insights into the organization of proteome, interactome and host protein recruitment. We envision that XL-MS will prove similarly useful for studying other viral particles, in particular of complex, large and enveloped viruses.

## MATERIAL AND METHODS

### Cells and viruses

Human embryonic lung fibroblasts (Fi301) were maintained as previously described ^49^. In preparation for SILAC proteomic analysis, cells were SILAC-labeled for at least five passages using lysine and arginine-deprived DMEM, supplemented with 10% dialyzed serum (cut-off: 10 kDa, PAN-Biotech), heavy (L-[^13^C_6_,^15^N_2_]-lysine (Lys8), L-[^13^C_6_,^15^N_4_]-arginine (Arg10)) or light L (natural lysine (Lys0) and arginine (Arg0)) amino acids. For phosphoproteome analysis we supplemented 200 mg/L L-proline. Labeling efficiency was checked using LC–MS/MS. Fi301 cells L were also used for preparation of viral stocks. The HCMV strain TB40-BAC4 ^50^ was used for all experiments. Infectious virus titers were determined by immunotitration and indicated as IE-forming units (IU) per mL, as previously described ^51^. Infection experiments were carried out using either a high (5 IU/cell) or low (0.05 IU/cell) multiplicity of infection (MOI). BAC mutations were created by traceless mutagenesis according to established protocols ^52^. Mutations were verified by Sanger sequencing and PCR. See Supplementary Table 6 for a list of mutagenesis primers.

### Immunoblotting

Gradient-purified virions were resuspended in PBS and adjusted to equal concentrations based on their optical density at 600 nm wavelength. Then, equal virion amounts were centrifuged for 60 min at 35,000 rpm, using a TLS-55 rotor (Beckman). The virions were lysed by sonication in 50 mM Tris-Cl (pH 6.8), 2% SDS, 10% glycerol, 100 mM dithiothreitol, 2 µg mL−1 aprotinin, 10 µg mL−1 leupeptin, 1 µM pepstatin, 0.1 mM Pefabloc, bromophenol blue and boiled at 95°C for 3 min. The samples were resolved by sodium dodecyl sulfate (SDS) polyacrylamide gel electrophoresis and blotted to polyvinylidene fluoride membranes. To prevent nonspecific binding, blots were incubated in Tris-buffered saline, 0.1% Tween-20, 5% skim milk. The following primary antibodies were used: anti-UL32 (clone XP1, provided by Bodo Plachter), anti-PP1 (clone E9, sc-7482, Santa Cruz), anti-14-3-3 (clone H8, sc-1657, Santa Cruz). Blots were developed using horseradish peroxidase-conjugated secondary antibodies in conjunction with suitable enhanced chemiluminescence detection systems.

### Flow Cytometry

Flow cytometric analysis of DNA content and IE1/IE2 expression was carried out as described previously ^49^. Alexa Fluor 488-conjugated anti-IE1/IE2 (clone 8B1.2, MAB810X, Merck) was used. Flow cytometry was performed using a FACSCanto II instrument and FACSDiva software (both from BD Biosciences). Cellular debris, cell doublets and aggregates were gated out of analysis.

### Virion purification and cross-linking

To prepare HCMV particles for *in situ* cross-linking, we first harvested infectious cell culture supernatants and clarified them from cellular debris by centrifugation at 1,500 rcf for 10 min. The remaining viral supernatant was centrifuged for 1 h at 25,000 rpm in an SW-28 rotor (Beckman). The resulting virus pellets were resuspended in 1-2 pellet volumes PBS and supplemented with 2.5 mM DSSO (100 mM stock solution in DMSO). The cross-linking reaction was incubated for 30 min at 25°C under shaking conditions (1000 rpm). The cross-linking step was repeated once with additional 2.5 mM DSSO before the reaction was quenched with 50 mM Tris-HCl (pH 8.0) for 20 min at 25°C under constant agitation (1000 rpm). Subsequently, the cross-linked material was loaded onto glycerol-tartrate gradients as described elsewhere ^53^ and centrifuged for 1 h at 25,000 rpm in an SW-40 rotor (Beckman), with brakes set at the slowest possible deceleration. The virion band was aspirated through the wall of the ultracentrifuge tube using a 1 ml insulin syringe equipped with a 1.2 x 40 mm needle. The virion fraction was washed twice in PBS. The first washing step included virion sedimentation at 30,000 rpm in an SW-60 rotor (Beckman), the second step sedimentation at 35,000 rpm in an TLS-55 rotor. The purified virions were stored at – 80°C for XL-MS sample preparation (see below).

### Virion sample preparation for XL-MS and bottom-up proteomics

First, to increase proteomic coverage of glycoproteins, cross-linked virion samples were deglycosylated using Protein Deglycosylation Mix II (P6044, NEB) under denaturing conditions according to manufacturer’s instructions. Lysis of virion samples was performed by adding 3 volumes of lysis buffer containing 8 M Urea, 1% Triton X-100, 30 mM chloroacetamide (CAA), 5 mM tris(2-carboxyethyl)phosphine hydrochloride (TCEP), 700 units Benzonase (70746, Merck) and incubation on ice for 30 min followed by sonication for 45 min (30 s on, 30 s off) in a Bioruptor Pico (Diagenode) at 4°C. Proteins were extracted using methanol-chloroform precipitation according to standard protocols ^54^, dried and resuspended in digestion buffer (50 mM triethylammonium bicarbonate (TEAB), pH 8.0, 1% sodium deoxycholate, 5 mM TCEP and 30 mM CAA). Proteins were digested by adding trypsin at an enzyme-to-protein ratio of 1:25 (w/w) and LysC at a 1:100 ratio (w/w) at 37°C overnight in the dark. Peptides from cross-linked samples were desalted using Sep-Pak C8 cartridges (Waters). Peptides from non cross-linked samples were desalted using C18 stage-tip purification followed by LC-MS.

Peptides destined for cross-link analysis were further fractionated by strong cation exchange using a PolySULFOETHYL A^TM^ column (PolyLC Inc.) on an Agilent 1260 Infinity II system. A 90 min gradient was applied and 33-35 fractions were collected, desalted by C8 stage-tips, dried under speed vacuum and subjected to LC-MS analysis.

### Cell sample preparation for LC-MS

Cells were harvested at 6 days post infection by scraping in PBS. Cell lysis, protein extraction and digestion were performed exactly as described above for virions.

### Affinity purification-MS (AP-MS)

PLFs were infected with HCMV-UL32-GFP (phosphatase-activity dependent interactome, GFP-AP), HCMV-UL32-HA or HCMV-UL32-SILK/RVxFmut-HA (PP1-binding dependent interactome, HA-AP) at an MOI of 5 IU/cell. At 5 days post infection, cells were treated with 300 nM calyculin A (BML-EI192-0100, Enzo Life Sciences) for 20 min or left untreated. Directly after, cells were harvested by scraping in PBS and processed as previously described ^55^. In brief, cells were L 1L MgCl_2_, 1% Nonidet P-40, 0.1% SDS, 5% glycerol, 1L DTT, 2 µg mM NaCl, mL^−1^ aprotinin, mM mM L L L mM Pefabloc, 0.5 mM Na_3_VO_4_, 10 mM β-glycerophosphate, 1 mM NaF). Wash and lysis buffer of calyculin A treated samples contained additional 20 nM Calyculin A. Lysates were sonicated to solubilize nucleocapsid-associated UL32-GFP before clearing the lysates for 20 min at 12.000 rcf at 4°C. For GFP-AP, GFP-trap agarose (gta-20, ChromoTek) was employed. Lysates were incubated with the agarose for 1 h, lysis buffer was used for the first two washing steps, lysis buffer without detergent for the third and final washing step. Proteins were eluted by incubating the beads in a total volume of 0.2 mL of 8 M L guanidine hydrochloride at 95°C under shaking. For HA-AP, a µMACS HA isolation kit (Miltenyi Biotec) was employed according to the manufacturer’s instructions, with the following modifications. Lysates were incubated with the the magnetic microbeads for 1L onto the microcolumns, lysis buffer was used for the first washing step, lysis buffer without detergent for the second and 25 mM Tris-HCl (pH 7.4) for the final washing step. Proteins were L eluted by adding 200 µl guanidine hydrochloride which had been prewarmed to 95°C.

Proteins were precipitated from the eluates by adding 1.8 mL LiChrosolv ethanol (Merck) and L 1LµL GlycoBlue (Thermo Fisher). After incubation at 4°C overnight, samples were centrifuged for 1 h at 4°C, ethanol was decanted and the pellet was air-dried before proteins were resolved in digestion buffer (see above), supplemented with trypsin at an 1:25 and LysC at an 1:100 enzyme-to-protein ratio (w/w). Digests were incubated overnight at 37°C, subjected to C18 stage-tip desalting followed by LC-MS analysis.

### Phosphopeptide enrichment

Peptides destined for phosphoproteome analysis were subjected to IMAC enrichment using a ProPac^TM^ IMAC-10 column (Thermo Fisher) on a 1260 Infinity II system (Agilent Technologies). A 30 min gradient was applied and the fraction corresponding to the phosphopeptides was collected, dried under speed vacuum and subjected to LC-MS analysis.

### LC-MS analysis

LC-MS analysis of cross-linked and SCX-fractionated peptides was performed using an UltiMate 3000 RSLC nano LC system coupled on-line to an Orbitrap Fusion Lumos mass spectrometer (Thermo Fisher). Reversed-phase separation was performed using a 50 cm analytical column (in-house packed with Poroshell 120 EC-C18, 2.7 µm, Agilent Technologies) with a 120 or 180 min gradient. Cross-link acquisition was performed using an LC-MS2 method. The following parameters were applied: MS resolution 120,000; MS2 resolution 60,000; charge state 4-8 enabled for MS2; stepped HCD energy 21, 27, 33.

LC-MS analysis of linear peptides (unmodified and phosphopeptides) was performed using a Dionex UltiMate 3000 system (Thermo Fisher) connected to a PepMap C-18 trap-column (0.075 x 50 mm, 3 μm particle size, 100 pore size; Thermo Fisher) and an in-house-packed C18 column μ (column material: Poroshell 120 EC-C18, 2.7 µm; Agilent Technologies) at 300 nL/min flow rate and 120-240 min gradient lengths. The MS1 scans were performed in the orbitrap using 120,000 resolution. Precursors were isolated with a 1.6 Da isolation window and fragmented by higher energy collision dissociation (HCD) with 30% normalized collision energy. The MS2 scans were acquired either in the ion trap or the orbitrap. For the ion trap, standard automatic gain control (AGC) target settings, an intensity threshold of 1e4 (5e3 for 240 min gradients) and maximum injection time of 50 ms were used. A 1 s cycle time was set between master scans. For MS2 acquisition in the orbitrap, we used standard AGC settings, an intensity threshold of 5e4, 50 ms maximum injection time and a resolution of 15,000 for unmodified peptides or 30,000 for phosphopeptides. A 2 s cycle time was set between master scans.

### XL-MS data analysis

Peak lists (.mgf files) were generated in Proteome Discoverer (version 2.1) to convert each .raw file into one .mgf file containing HCD-MS2 data. The .mgf files were used as input to identify cross-linked peptides with a stand-alone search engine based on XlinkX v2.0 ^56^. The following settings of XlinkX were used: MS ion mass tolerance, 10 parts per million (ppm); MS2 ion mass tolerance, 20 ppm; fixed modification, Cys carbamidomethylation; variable modification, Met oxidation; enzymatic digestion, trypsin; and allowed number of missed cleavages, 3; DSSO cross-linker, 158.0038 Da (short arm, 54.0106 Da, long arm, 85.9824 Da).

All MS2 spectra were searched against concatenated target-decoy databases generated based on the virion proteome determined by bottom-up proteomics, containing 1,318 target sequence entries. Raw-files from both biological replicates were searched combined and cross-links were reported at 1% FDR at unique lysine-lysine connection level based on a target-decoy calculation strategy using randomized decoys. All identified cross-links can be accessed in Supplementary Table 1. Quality control analyses (related to Fig.s 1C-E), clustering of viral proteins (related to Fig. 2A), analysis of scaffold indices (related to Fig. 2B) and analysis of domain-specific interactions to UL32 (related to Fig.s 2 C-D) were performed with this set of cross-links. All other analyses are based on the filtered set of cross-links (see below), representing a high confidence PPI-map of the particle.

For this filtering, it was first checked whether cross-links were identified in both replicates independently at 1% FDR. We required that cross-links from abundant virion proteins (> 10 copies) be identified in both replicates. When a cross-link involved a lower abundant protein (that is, < 10 copies) it was required that the cross-link is only identified in one of the replicates. The rationale is that this balances cross-link confidence for highly abundant proteins with sensitivity for low abundant proteins. Second, PPIs were only reported when there were two unique cross-links identified. Third, host proteins were removed that did not directly link to viral proteins. The filtered set of cross-links is available in Supplementary Table 2. The XL-based PPI network was generated on the basis of inter-protein cross-links from Supplementary Table 2 and visualized using edge-weighted spring-embedded layout ^57^ in Cytoscape v3.7.2. Edge weighting is based on the number of identified inter-links between protein pairs. Additional cross-link networks between selected proteins were visualized using xiNET ^58^.

For clustering of viral proteins across virion (sub)layers, we used the full set of identified cross-links at 1% FDR (Supplementary Table 1). Only cross-links of viral proteins (intra-links and inter-links) were considered and viral proteins were excluded when (i) less than 10 of such cross-links were identified or (ii) the respective protein only formed intra-links. The cross-link count between any protein pair (“interactor 1” and “interactor 2”) was then divided to the total of cross-links for one of the proteins (“interactor 1”) to yield PPI specificity values. Correlation-based clustering was performed (1-Pearson’s R) on the resulting matrix using complete linkage and data was visualized in R (v4.1.2.)/Rstudio (v.1.3.1093) using heatmap.2 function from gplots package. For calculating the scaffold index, we removed the PPI specificity values for interactions linking the same protein and summed up the PPI specificity values for each PPI of the interactor 2 proteins within the sub-layers.

The circular heatmap was generated by circos.heatmap function in circlize R package and clusters are based on euclidean distances. Cross-links of host proteins to viral proteins of the specific clusters (as obtained from the analysis in Fig. 2A, exempt UL48A, which we considered as nucleocapsid ^8^), were z-scored by subtracting the mean and dividing by the standard deviation.

### Bottom-up proteomics data analysis

Raw data were analyzed and processed using MaxQuant 1.6.2.6 software ^59^. Search parameters included two missed cleavage sites, fixed cysteine carbamidomethyl modification, and variable modifications including methionine oxidation, N-terminal protein acetylation. In addition, serine, threonine, and tyrosine phosphorylations were searched as variable modifications for phosphoproteome analysis. Arg10 and Lys8 were set as labels where appropriate (double SILAC samples). The “second peptide” option was enabled and peptide mass tolerance was 6 ppm for L MS scans and 20Lppm for MS/MS scans. “re-quantify”, “iBAQ” and “LFQ” were enabled where appropriate. Database search was performed using Andromeda, the integrated MaxQuant search engine, against a protein database of HCMV strain TB40-BAC4 ^50^ and a Uniprot database of homo sapiens proteins (downloaded 2020) with common contaminants. FDR was estimated based on target-decoy competition and set to 1% at peptide spectrum match (PSM), protein and modification site level. For subsequent analysis, we used proteinGroups.txt, peptides.txt or Phospho (STY).txt (phosphoproteomics) MaxQuant output files with potential contaminants, reverse database hits and proteins only identified by (modification) site removed.

### Bottom-up proteomics data processing

For label-free AP-MS data, LFQ values from proteinGroups.txt were log2 transformed and two-sample t-test p-values between experimental groups were calculated along with the average fold-change difference.

For SILAC-based quantification of virion-level protein differences, we used SILAC ratios as normalized by MaxQuant. For the corresponding phosphoproteome data, only sites with a localization probability > 0.75 were considered (from Phospho (STY).txt table). The position of individual phosphosites as identified from this experiment were used to map phosphosites on the UL32 amino acid sequence (Fig. 4C). Phosphosite SILAC ratios were also corrected for the protein-level SILAC ratios (from proteinGroups.txt) as quantified from an analysis of the whole virion proteome without IMAC-enrichment. Phosphosites were excluded when the corresponding phosphopeptide contained residues mutated in the SILK/RVxF motifs of UL32. Then, protein-level corrected phosphosite ratios were averaged (quantification in both replicates required).

For comparing phosphosite and peptide abundances in label-free AP-MS samples, Maxquant output tables Phospho (STY).txt and peptides.txt were used, respectively. Phosphosite and peptide intensities for UL32 were summed up across the three replicates.

Enrichment levels comparing virions versus cells were calculated based on log2 transformed LFQ intensities in MaxQuant output file proteinGroups.txt using perseus software ^60^. Therefore, the proteinGroups file was filtered by requiring at least four values in either virion or cell samples. Missing values were imputed based on a normal distribution shrinked by a factor of 0.3 and down-shifted by 1.8 standard deviations. T-tests, log2 differences, and spearman’s correlation coefficients (rho) were calculated based on these values.

### Absolute quantification of protein copy numbers

We absolutely quantified the copy numbers of host and viral proteins in purified particles based on intensity based absolute quantification (iBAQ)-values ^61^. As calibrators, we used proteins with known copy numbers, associated with the nucleocapsid ^8^ and portal complex ^11^. Therefore, iBAQ-values from MaxQuant output file proteinGroups.txt were extracted. We then used the following known copy numbers of HCMV proteins as calibrators for a linear regression analysis: UL93, UL104: 12 copies; UL77: 24 copies; UL46: 320 copies; UL85: 640 copies; UL86, UL48A, UL32: 960 copies ^8, 11^. Slope and offset were used to calculate unknown copy numbers of host and viral proteins according to Eq. 1.

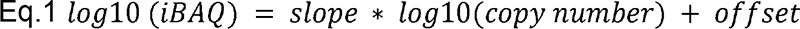

Total protein copies were summed up for all identified proteins and divided by the volume of a sphere of 200 nM diameter to yield protein density. Protein mass was calculated by multiplying the total protein copy number with the average mass of a human protein (44 kDa), as previously calculated ^36^.

### 14-3-3 binding site prediction

We used the freely available 14-3-3pred tool (https://www.compbio.dundee.ac.uk/1433pred) ^37^ to map putative 14-3-3 binding sites. Serines or threonines with a consensus score > 0.6 were considered.

### Structure mapping and calculations

Mapping of cross-links to cryoEM structures of UL55 (pdb 7kdp, 7kdd) was done using the PyMOL Molecular Graphics System, Version 2.0 Schrödinger, LLC. The shortest possible distances between individual chains were considered. One cross-link with distance greater than 40 L in pre-fusion conformation could not be mapped to the post-fusion structural model due to missing coordinates at Lysine residue 700.

Electrostatic surface charges of a nucleocapsid hexon (pdb 5vku) were calculated using the APBS Tools 2.1. plugin to pymol and individual cross-linked lysines were highlighted as spheres.

The length of the UL32 C-terminus was estimated based on the contour length of an amino acid of 3.4 L ^62^ (BNID:114332) and multiplied by 763 amino acids.

### Electron microscopy

Human fibroblasts infected with HCMV-UL32-GFP were fixed with 4% formaldehyde for 120 min while shaking. After fixation, cells were pelleted, cryoprotected in 2.3 M sucrose and plunge-frozen on pins for Tokuyasu sectioning ^63^. For immunogold labeling, ultrathin sections were collected on coated grids, blocked and co-stained by anti-GFP (1:250, 132005, Synaptic Systems), anti-pan 14-3-3 (1:50, clone H8, sc-1657, Santa Cruz) and anti-PP1 (1:50, anti-PP1, clone E9, sc-7482, Santa Cruz) followed by secondary antibodies gp-12 nm gold (species: guinea pig) and ms-18 nm gold (species: mouse). After washing, sections were contrasted and covered by polyvinyl alcohol and tungsto-silicic acid hydrate. Stained ultrathin sections were examined with a Zeiss 902 kV and photographs were taken with a Morada G2 TEM-L Camera.

## ACKNOWLEDGMENTS

The authors thank Dmytro Puchkov for valuable advice and interpretation regarding electron microscopy experiments and Heike Stephanowitz for excellent technical assistance. The authors thank Barth van Rossum for fruitful discussion on the possible structural arrangement of UL32. BB was funded from DFG grant BO 5917/1-1 and Leibniz-Wettbewerb P70/2018.

## AUTHOR CONTRIBUTIONS

Conceptualization: BB, LW, FL; Methodology: BB, IG, SH, LW, FL; Investigation: BB, IG, LM, JP, SH, LW; Formal Analysis: BB, IG, LM, JP, SH, LW; Resources: IG; BV; LW; FL; Visualization: BB, JP; Data Curation: BB; Funding acquisition: BB, LW, FL; Project administration: BB, LW, FL; Supervision: ML, LW, FL; Writing – original draft: BB; Writing – review & editing: BB, LW, FL.

## DATA AVAILABILITY

The mass spectrometry proteomics data have been deposited to the ProteomeXchange Consortium via the PRIDE ^64^ partner repository with the dataset identifier PXD031911. Reviewer account details: **Username:**; **Password:** n6JebLCu.

## Fig. LEGENDS

***Extended Data Fig.1. Data filtering and reproducibility.***

**(a,b)** Histogram of distances between C α atoms obtained from mapping cross-links onto the cryoEM structural models of homotrimeric UL55 in pre- **(a)** or post-fusion **(b)** conformations (pdb: 7kdp, 7kdd). **(c)** Direct comparison of distances between pre- and post-fusion conformations of UL55. **(d)** The number of cross-links is shown that survive the indicated filtering steps. **(e)** Venn diagrams comparing the number of PPIs between both replicates in unfiltered (Supplementary Table 1) and filtered (Supplementary Table 2) dataset. Homomeric interactions (based on intra-links) were removed for this analysis.

***Extended Data Fig.2. Host protein recruitment to HCMV particles.***

**(a,b)** Host proteins included in the PPI map (see Figure 1f) were compared to host proteins from individual **(a)** or **(b)** combined published proteomics dataset of purified HCMV particles. **(c)** Cross-link network between ribosomal 40S proteins and UL86 protein. **(d)** Cross-linked lysines were mapped onto a nucleocapsid hexon (UL86 hexamer), depicted from top (left side) (i.e. inner tegument side) or bottom (right side) (i.e. DNA-accommodating side). Electrostatic surface rendering with charges depicted ranging from negatively charged (red) to positively charged (blue) (pdb 5vku). **(e)** Depiction of cross-links between glycoprotein UL55 and tetraspanins CD9 and CD63. Domain annotations were obtained from uniprot.org.

***Extended Data Fig. 3. Quantitative proteomics identifies selective recruitment of host proteins.***

**(a)** Experimental setup to identify selectively recruited proteins (n=4 biological replicates). **(b)** Spearman’s correlation coefficients comparing log2 transformed LFQ-values of the experiment outlined in a. **(c)** Cross-comparison of protein copy numbers and enrichment levels of host and viral proteins. Selected proteins are depicted with their gene symbols. Enrichment levels are based on the mean of n=4 biological replicates and copy numbers based on the mean of n=2 biological replicates (with technical duplicates or triplicates). **(d,e)** Cross-link networks between selected host-virus PPIs.

***Extended Data Fig.4. Characterization of the 14-3-3 binding-deficient viral mutants.***

**(a,b)** EM images of the intracellular environment from HCMV-UL32-GFP infected PLFs (MOI=5) prepared using Tokoyasu method. Ultrathin sections were stained with immunogold against GFP (12 nm gold) and pan 14-3-3 or PP1 (18 nm gold), as indicated. Scale bars: 100 nM. Magnified views of the boxed regions in white are depicted in Fig 4.b. **(c)** Experimental workflow for SILAC-based comparison of virion protein content between mutant and WT viruses. The experiments were performed in label-swap duplicates. **(d)** Purified virions from WT or mutant viruses harboring alanine substitutions in the 3/4/5 region (see also Fig. 4c) were assessed for UL32, PP1 and 14-3-3 levels by immunoblotting. Control experiment to panel (e). **(e)** SILAC-based proteomic comparison of log2 protein fold-changes comparing purified particles of WT and 14-3-3 binding mutants for both replicates, based on n=2 biological replicates.

***Extended Data Fig.5. Characterization of the PP1 binding-deficient viral mutant.***

**(a)** Crystal structure of PP1 (pdb: 1fjm) with highlighted SILK and RVxF binding grooves as well as lysines cross-linked to UL32. **(b)** SILAC-based proteomic comparison of protein abundance comparing purified particles of wild-type and PP1-binding mutant viruses. **(c)** Cross-links between DYRK1A kinase and UL32. Cross-link reactive lysines are highlighted blue in the sequence of both UL32 and DYRK1A. **(d)** Average log2 phosphosite fold-changes comparing wild-type and PP1-binding mutant purified particles. Phosphosites belonging to host proteins, viral proteins (excluding UL32) or to UL32 were compared using a two-sided wilcoxon rank sum test. The average phosphosite ratio corrected by the protein ratio of n=2 biological replicates (including label-swap) are depicted. Experimental design as in Extended Data Fig.5C with additional phosphopeptide enrichment.

***Extended Data Fig.6. PP1-UL32 binding promotes viral growth and the onset of viral gene expression.***

**(a)** Growth curve of mutant and wildtype viruses on primary lung fibroblasts (PLFs) (MOI=5). Means (center of the error bars) and standard deviations of the mean of *n*L= 3 biological L replicates are depicted. Unpaired two-sided *t*-tests without multiple hypothesis correction were performed comparing the indicated viruses and time points. **(b)** Flow cytometry analysis of Immediate Early (IE) protein levels as a function of the cell cycle stage. PLFs were infected with the indicated recombinant viruses (MOI = 5, 6 h post infection). The percentage of IE1/2-positive cells in G1 or S/G2 compartments is given

***Extended Data Fig.7. Recruitment of PP1 functionally antagonizes 14-3-3 binding to UL32.***

**(a,c)** Experimental strategy to quantify differences in the PP1-binding **(a)** and phosphatase activity dependent-interactome of UL32 **(c)**. Both experiments were performed in n=3 biological triplicates. See also Fig. 5 c,d. **(b,d)** Volcano plot analysis showing p values based on a test of n=3 biological replicates and the average log2 fold-change. Cut-offs were set manually at P = 0.01 and log2 fold-change = 1 in (b) and log2 fold-change = 2 in (d). Proteins passing these cut-offs were labeled with gene symbols.

