## Supplementary figures and images for "Spatial, Quantitative and Functional Deconstruction of Virus and Host Protein Interactions Inside Intact Cytomegalovirus Particles"

### Extended Data Fig. 1

#### Extended Data Figure 1

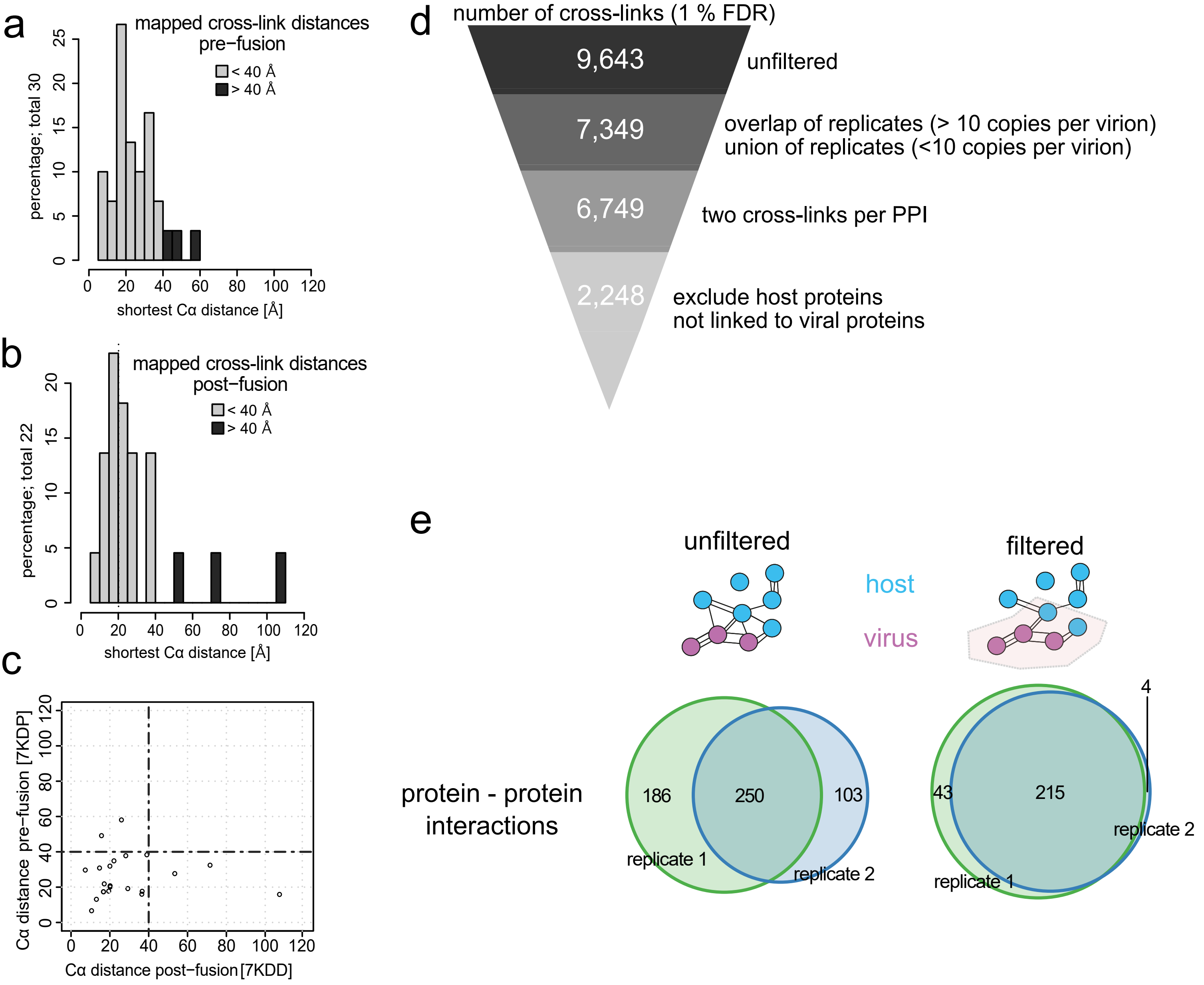

### Extended Data Fig. 2

#### Extended Data Figure 2

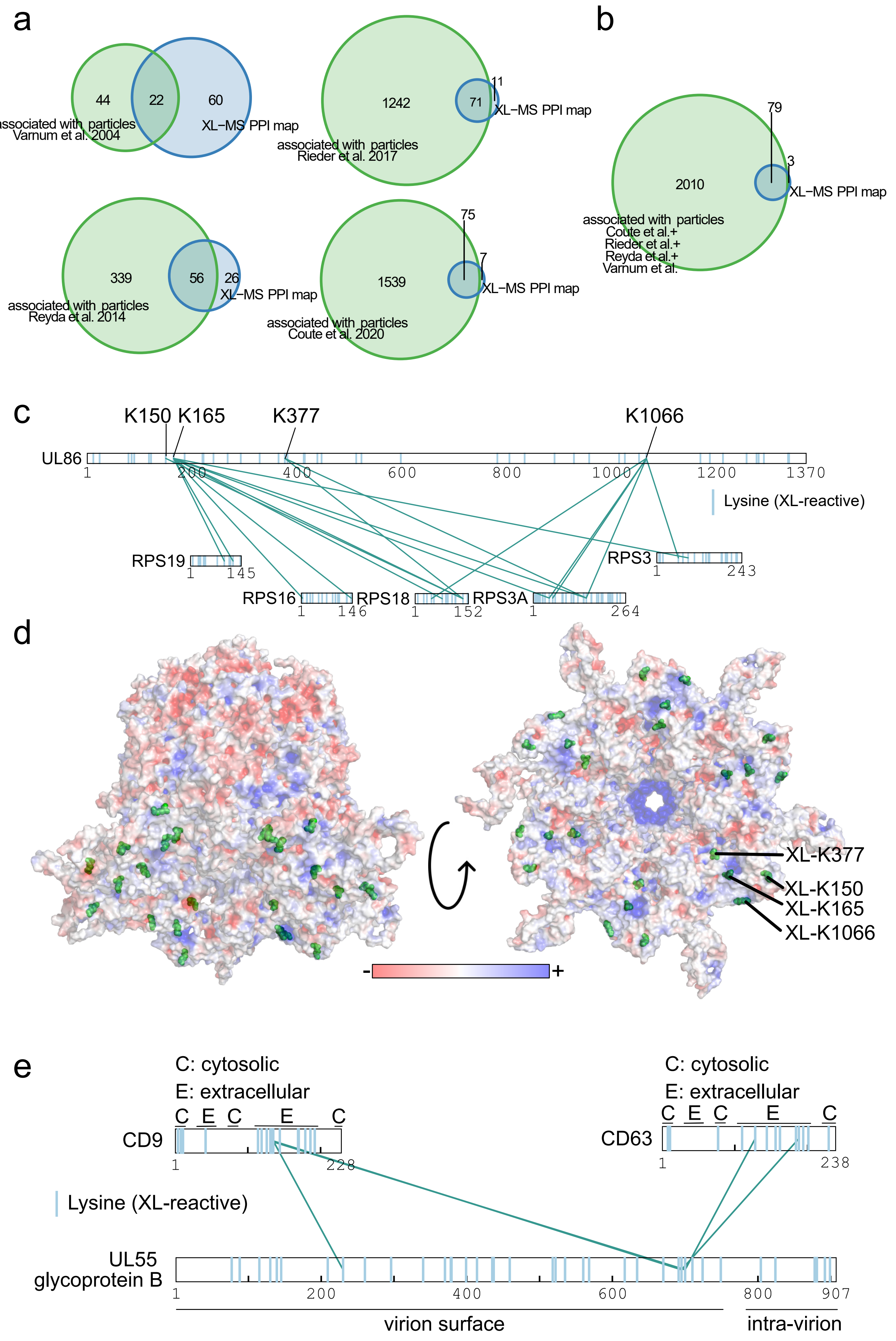

### Extended Data Fig. 3

#### Extended Data Figure 3

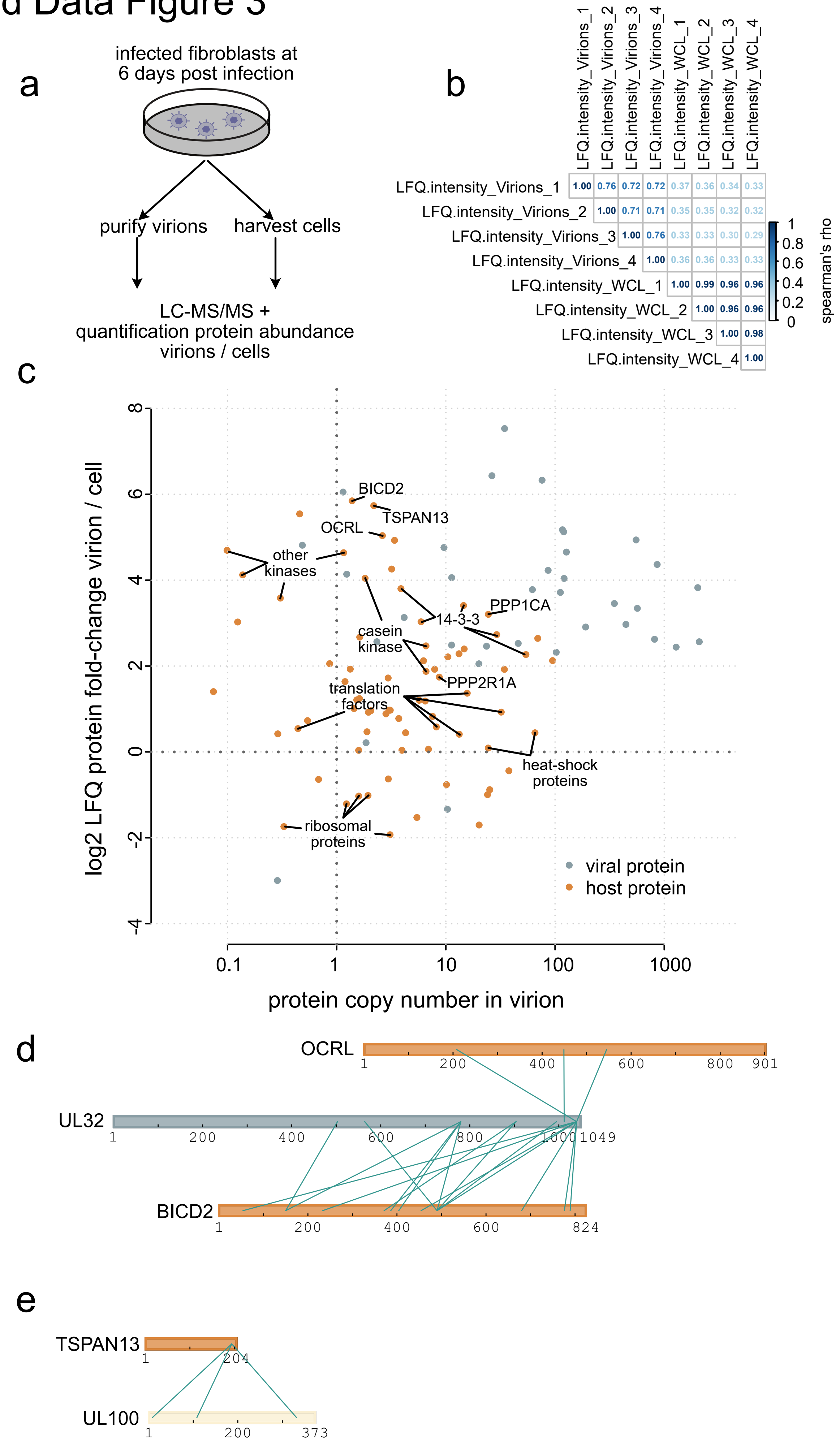

### Extended Data Fig. 4

#### Extended Data Figure 4

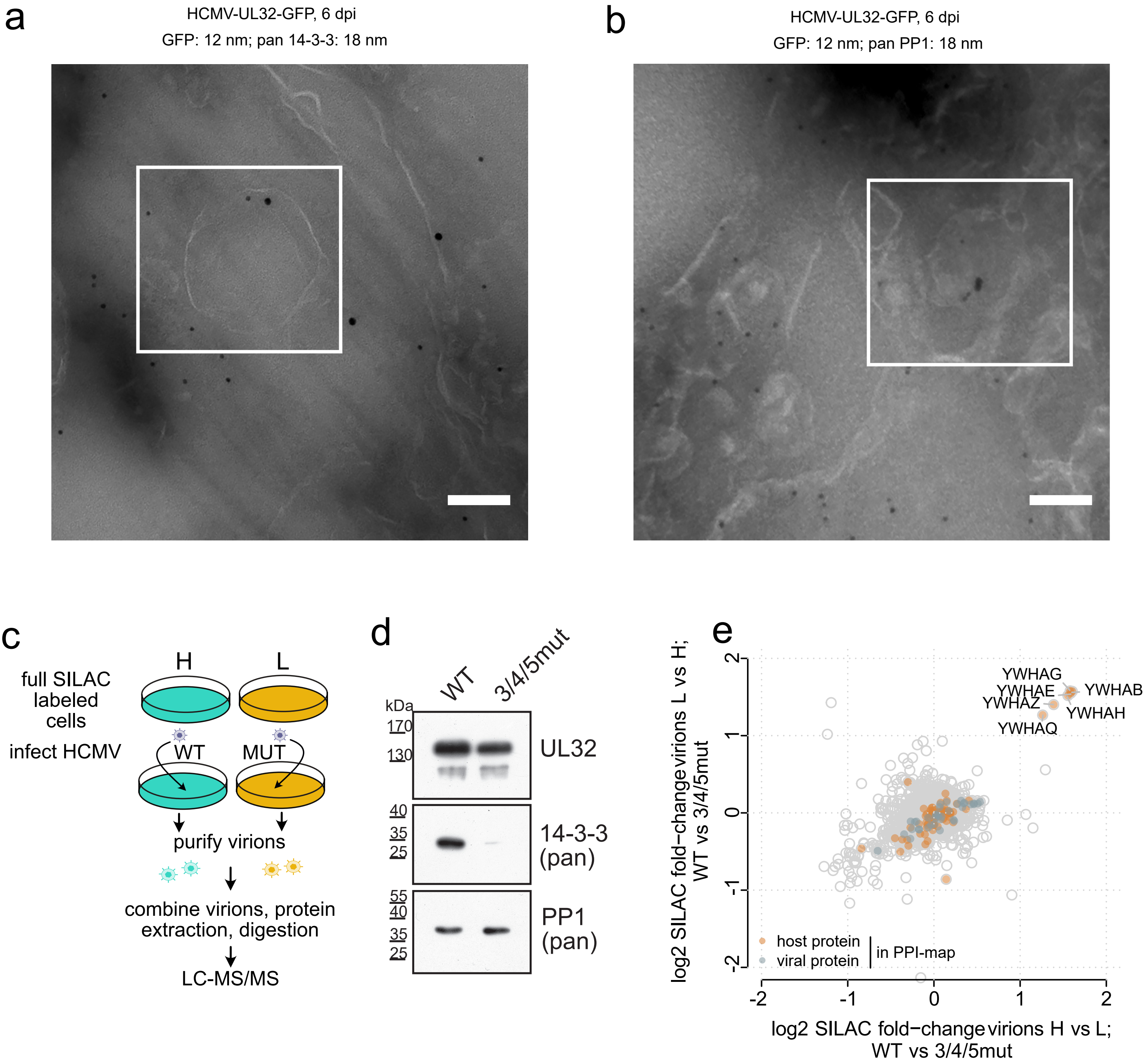

### Extended Data Fig. 5

#### Extended Data Figure 5

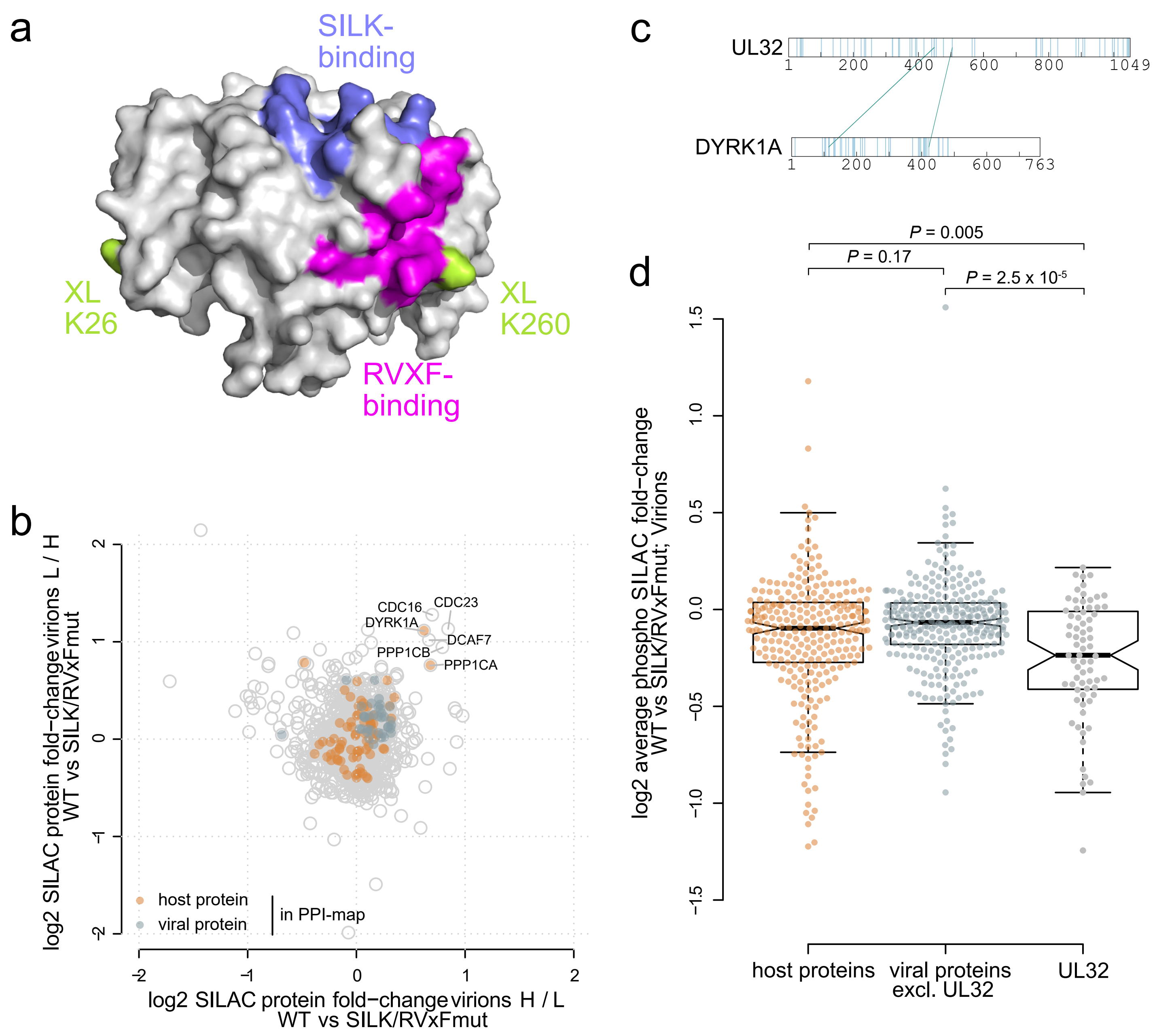

### Extended Data Fig. 6

#### Extended Data Figure 6

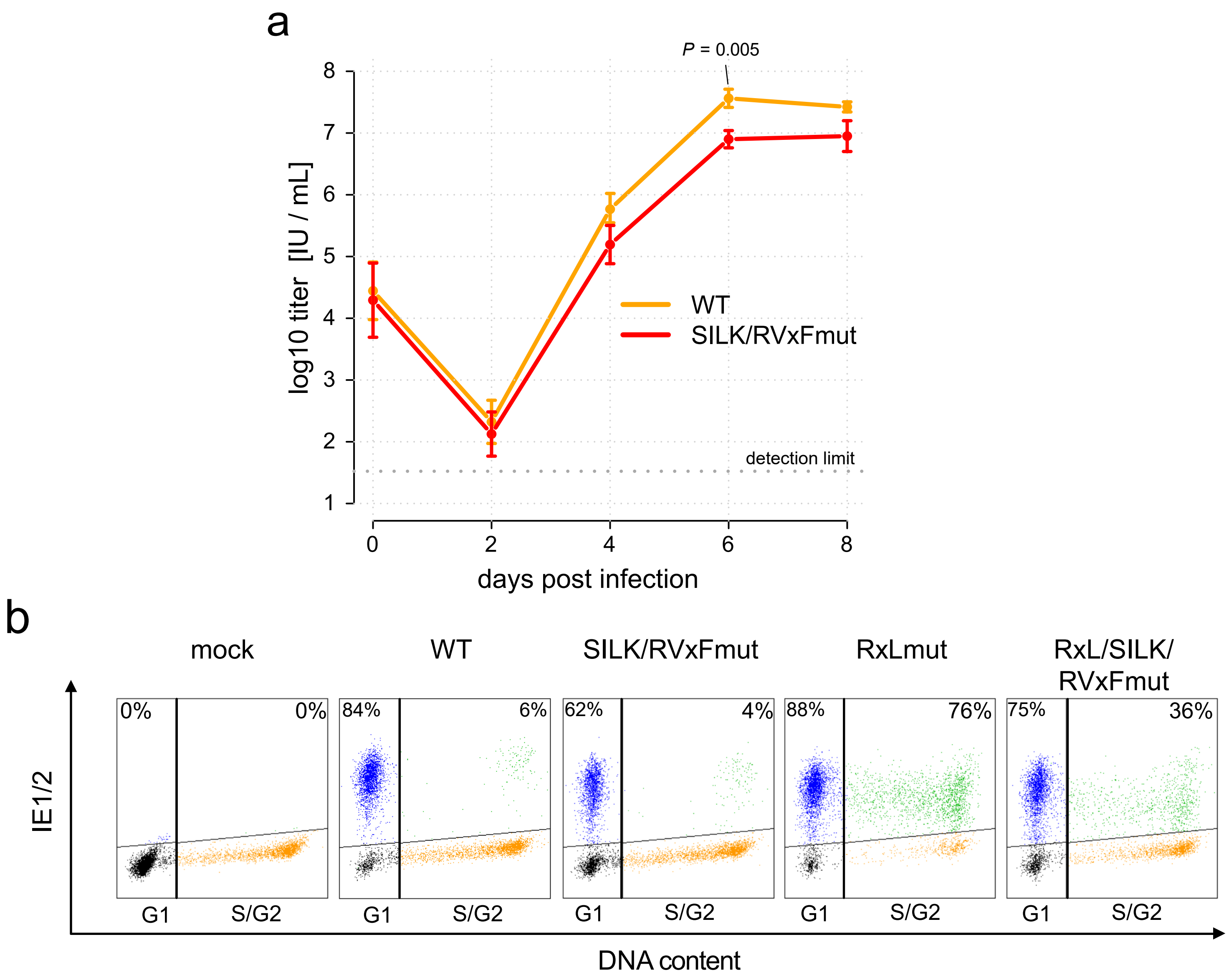

### Extended Data Fig. 7

#### Extended Data Figure 7

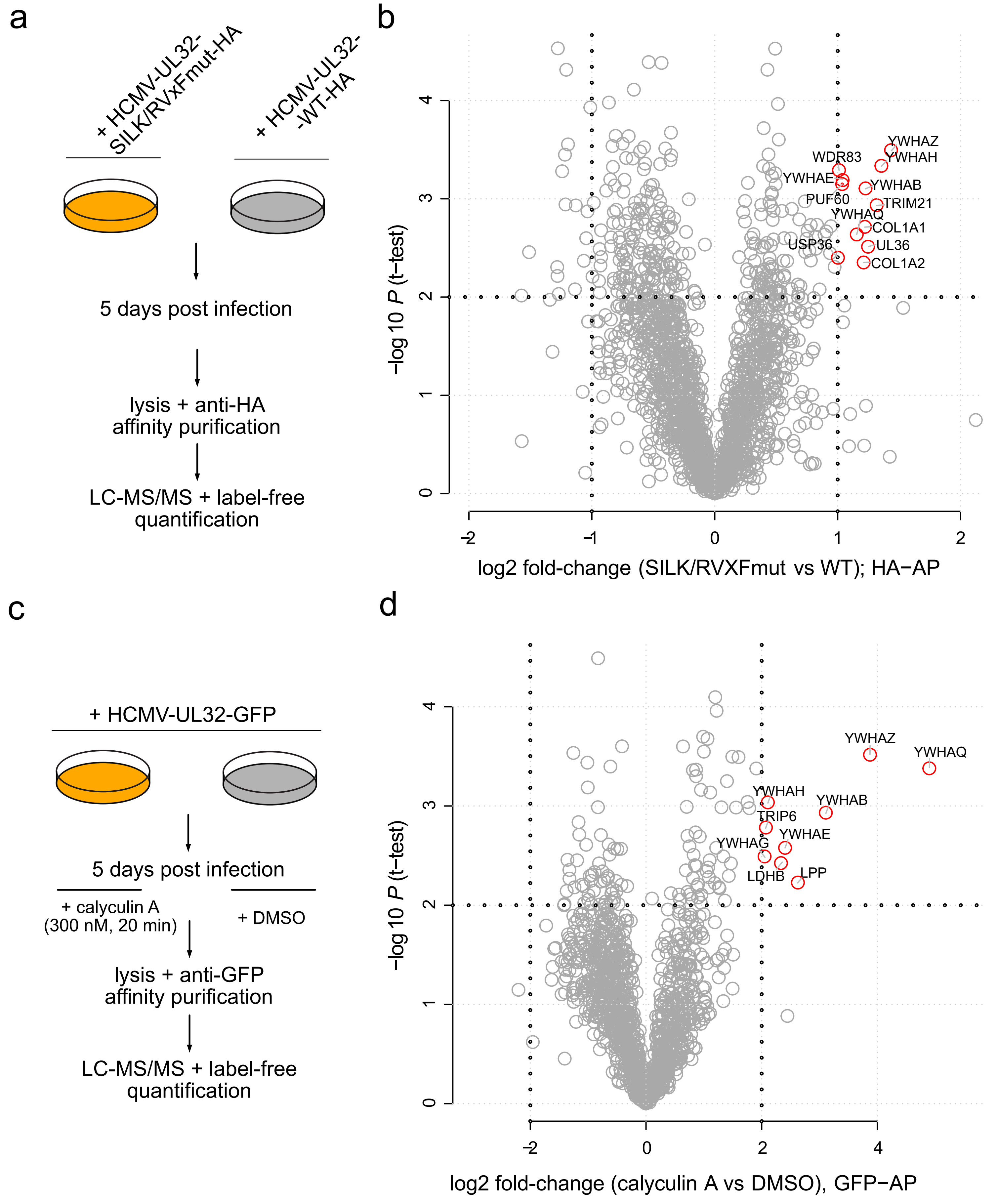
